# *Ardisia mela* (Primulaceae) a new threatened, herbaceous species of *Ardisia* subgen. *Kamardisia* from the forest of the Crystal Mts, Gabon

**DOI:** 10.64898/2026.07.29.740512

**Authors:** Harry Murdoch, Martin Cheek

## Abstract

*Ardisia mela* is formally described from the forests of the Crystal Mts, Gabon based on a specimen collected in 1968 and informally described in 1979 by Taton as Species 2. Despite the proximity of the collection to a road, and a resurgence of botanical surveys in the Crystal Mts in recent years, the species appears not to have been refound, and as it occurs in a logging concession and areas of cleared forest can be seen in the vicinity of the collection site, the species is considered threatened and provisionally assessed as Critically Endangered using the 2012 IUCN standard.

## Introduction

The nine herbaceous species of *Ardisia* Sw. in continental Africa were delimited, recognised as subgenus *Kamardisia* Cheek and revised in Peng & Cheek (2025). An introduction to the taxonomy, chemistry and uses of the genus, and the discovery of species in Africa can be found in in Peng & Cheek (2025). In this paper we described a tenth species, overlooked in the work referred to, but informally described as *Ardisia* sp 2 (Taton 1979). This taxon had been included in *A. ebo* Cheek (Cheek and Xanthos 2012) on the basis of the description only. However, this error was discovered on visiting P in July 2025, since on viewing the specimens there is no confusing the two species, *A. ebo* having elliptic, very thin leaves, with an acute apex and base and a crenate margin, while *Ardisia* sp 2 has thick, obovate leaves, with a rounded apex and obtuse base, and an entire margin. In this paper we formally describe *Ardisia* sp 2 as *A. mela* and give it a provisional conservation assessment. We also compare it with the similar *A. sadebeckiana* Gilg, and we revise and update the key to the herbaceous African *Ardisia* species (Peng & Cheek 2025) to accommodate the new species.

This paper is one of several preparatory to the Flore Du Gabon account of *Ardisia*. It concludes descriptions of the species of the herbaceous chamaephyte group (see African *Ardisia* life form classification, Cheek *et al*. 2025) and follows revision of the monocaul phanerophyte species (Murdoch & Cheek 2026).

## Materials and Methods

See Peng & Cheek (2025).

### Taxonomic Treatment

#### Key to the herbaceous species of *Ardisia* subg. *Kamardisa* in continental Africa (updated from Peng & Cheek 2025)

1. Leaves with red, circular or slightly lobed, peltate scales conspicuous on at least the abaxial leaf-blade (c. 0.1 mm diam., visible with a 10× hand lens)____________________________________________________6 – Leaves lacking circular, peltate scales on the abaxial blade surface (< 0.1 mm diam., not visible with a 10× hand lens)____________________________________2
2. Leaves 3 – 7(– 8) cm wide; flowers 3 – 6 per inflorescence; calyx lobes >1 mm long___________3 – Leaves (1.5 –) 1.9 – 2.8 cm wide; flowers 2(– 3) per inflorescence; calyx lobes 0.7 – 0.8 mm long____________________________________________________________***A. ebo***
3. Leaf-blade 5.5 – 7(– 8) cm wide; obovate, base cordate, margin entire_______________4 – Leaf-blade 3 – 5 cm wide; elliptic, base acute, margin crenate___________________________5
4. Leaf-blade base apex mucronate, base cordate; petiole1.2 – 2 cm long; calyx c. 2 mm long; pedicels 8 – 10 mm long________________________________________***A. sadebeckiana*** – Leaf-blade base apex rounded, base obtuse; petiole 0.5 – 1.3 cm long; calyx c. 1 mm long; pedicels c.4 mm long ***A. mela***
5. Petiole 7 – 10(– 18) mm long; leaf-blade discolorous, dring black above, dull white below_________________________________________***A. schlechteri*** – Petiole 3 – 5(– 10) mm long; leaf-blade ±concolorous, drying green on both surfaces***__________________________________________A. massaha***
6. Leaves 4.4 – 5.3 × 2.1 – 2.2 cm, apex acuminate________________________________***A. minuta*** – Leaves 4.3 – 9.3(– 12.2) × (2.7 –)5 – 6.3 cm, apex acute or rounded 7
7. Leaf apex broadly rounded, base abruptly cordate_______________________________________***A. ngounie*** – Leaf apex acute, base obtuse or rounded 8
8. Leaves rhombic, the distal ½ to ⅔ with distinct marginal teeth; pedicels glabrous apart from sessile globose, red glands; fruit with 5 longitudinal lines; leaf nervation inconspicuous_____________________***A. chaillu*** – Leaves elliptic or ovate-lanceolate, margins entire or sinuous, sometimes with inconspicuous marginal teeth proximally and distally; pedicels with broad, short acute hairs, non-glandular; fruit (where known) xlacking longitudinal lines; leaf nervation conspicuous 9
9. Peltate scales present on adaxial as well as abaxial blade surface; leaf length: width ratio <2; secondary and tertiary veins raised, conspicuous on adaxial surface of leaf; calyx lobes about as long as wide_________________________________________________________________________***A. hansii*** – Peltate scales present only on abaxial blade surface; leaf length: width ratio>2; secondary and tertiary veins inconspicuous on adaxial surface of leaf; calyx lobes nearly twice as broad as long_______________________________________________________***A. waka***

***Ardisia mela*** *Cheek & Murdoch* **sp. nov**. Type: Gabon, Estuaire province, Crystal Mountains, 10 km sud Mèla (10 km S of Mèla), 0°36’ N, 10°18’ E, fr. 2 Feb. 1968, *Hallé & J*.*F. Villiers* 4877 (holotype P barcode P00443366!; isotype P0044336671!).

*Ardisia* sp. 2 (Taton 1979: 118).

*Ardisia ebo* sensu Cheek & Xanthos (2012: 281) in part: *Hallé & J*.*F. Villiers* 4877!

*Small creeping herb* c. 15 cm tall. Stems decumbent, creeping horizontally, terete, 2.5 – 3 mm diam.; distal part unbranched, producing adventitious roots up to 14 cm long, arising up to 5 cm above the ground on the vertical stem; new stems arising 1 – 10 cm apart along the creeping portion, internodes 8 – 13 mm long, phyllotaxy distichous, indumentum rusty brown tomentellous, becoming sparser proximally, epidermis red-brown. *Leaves* alternate, petiole canaliculate, 5 – 13 × 1.5 – 2 mm; lamina discolorous, adaxially dark brown, abaxially paler orange-brown, obovate to suborbicular, 5.4 – 10 × 4.6 – 6.6 cm, length to width ratio 1.1 – 1.5:1, apex rounded, base obtuse, margin entire, midrib conspicuous, raised on the adaxial and abaxial surfaces, grooved on adaxial; lateral nerves 4 – 8 on each side of the midrib, departing from midrib at 55 – 80º, brochidodromous, looping 2 – 4 mm from margin, tertiary nerves inconspicuous in reflected light, conspicuous in transmitted light, reticulate creating areolae; oil glands orange-red (transmitted light), brown to black in reflected light, visible and raised on adaxial and abaxial surface in reflected light (lens needed), suborbicular, elliptic or oblong, 0.09 – 1.06 × 0.09 – 0.29 mm, 3 – 11 per 2 × 2 mm; 1 – 11 glands per areole, scales conspicuous (microscope), abaxially red-brown, 23 – 47 per 2 × 2 mm, 0.04 – 0.06 mm wide, adaxially adpressed, red-brown, 14 – 33 per 2 × 2 mm, 0.04 – 0.06 mm wide. *Inflorescences* axillary, c. 6 per stem, c. 3-flowered, sessile, rachides 1 – 1.5 mm long, pedicel bases exserted from the rachis, rusty brown floccose hairs on rachides and bracts; bracts caducous, ovate-triangular, c. 2 × c. 1 mm; pedicels terete, broadening distally, c. 4 × c 0.75 mm. *Flower buds* and *Flowers* not seen. *Fruits* globose, maroon-brown, c. 6 mm diam. Calyx c. 1 mm long, cupular at base, lobes 5, reflexed, ovate-triangular, apex acute to subrounded, oil glands either orange or dark brown-black, subcircular to oblong. Fruit wall c. 0.06 mm thick; surface c. 20% covered by oil glands; oil glands black to dark reddish brown, subcircular, elliptic or oblong, 0.12 – 0.48 × 0.12 – 0.19 mm. Fig.1.

**Fig. 1.**
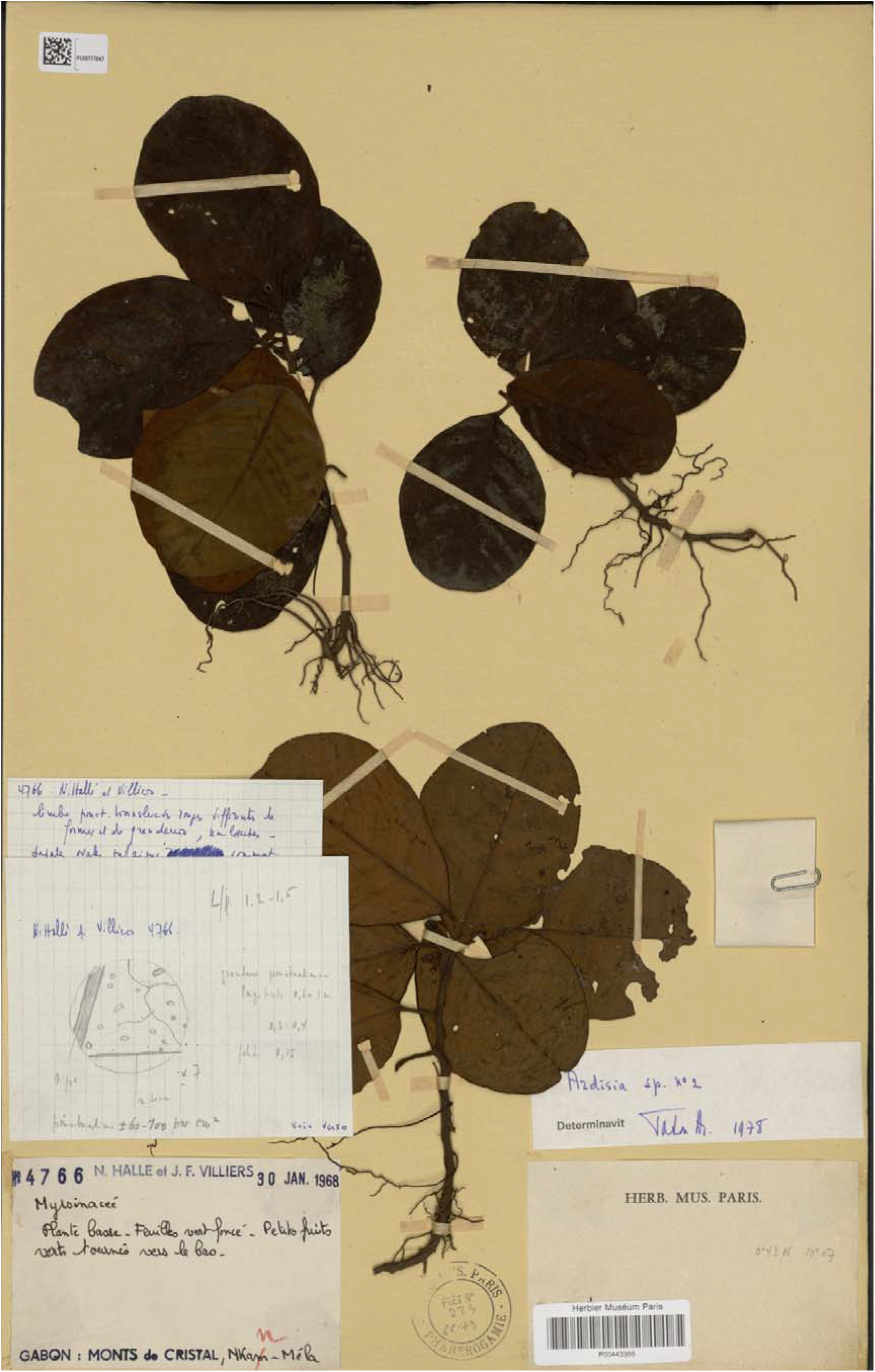
*Ardisia mela Hallé & J.F. Villiers* 4877 (P) holotype and only known specimen.

##### DIAGNOSIS

In the key to the herbaceous species of African *Ardisia* Subg. *Kamardisia* (Peng & Cheek 2025), because the abaxial leaf peltate scales are <0.1 mm diam. (not visible with a hand lens), the leaves 4.6 – 6.6 cm wide, flowers 3 per inflorescence, this taxon keys to couplet 3, and *A. sadebeckiana* Gilg. It differs from this species in the leaf-blade base apex rounded, the base obtuse, the petiole 0.5 – 1.3 cm long (vs mucronate, cordate, petioles 1.2 – 2 cm), it also differs in the calyx c. 1 mm long and pedicels c. 4 mm long (vs calyx c. 2 mm long; pedicels 8 – 10 mm long).

##### DISTRIBUTION

*Ardisia mela* is endemic to the Crystal Mountains, northeast of Libreville.

##### SPECIMENS EXAMINED. GABON

Estuaire province, Crystal Mountains, 10 km sud Mèla (10 km S of Mèla), 0°36’ N, 10°18’ E, fr. 2 Feb. 1968, *Hallé & J*.*F. Villiers* 4877 (holotype P! Barcode P00443371).

##### HABITAT & ECOLOGY

Estimated altitude of 600 m. Ecology unknown, satellite imagery shows dense evergreen forest.

##### CONSERVATION STATUS

Despite searches of gbif.org and of all Gabonese material of the genus at BM, K, P and WAG, *Ardisia mela* is known from only one collection in Gabon’s Crystal Mountains, 10 km south of Mèla (also known as Mala), a small settlement c. 90 km from the capital, Libreville. This collection is not from an actual or proposed protected area Texier *et al*. (2024) and exists within a logging concession. Logging represents a significant threat to this species since even in low density logging, habitat loss results from the creation of loading bays and logging roads, and canopy opening can be negative for herb species adapted to deep forest shade e.g. causing leaf bleaching. The georeference is not accurate since it predates GPS availability and was likely read from a map. Within 4 km of the point given on the label, areas of forest in the process of being cleared can be seen, with fallen tree trunks. The species has not been collected since 1968 despite its range being frequently surveyed, so the species may already be extinct. Following IUCN (2012, 2024), the area of occurrence (AOO) of this species is estimated as 4 km^2^, below the upper threshold for “Critically Endangered” status under Criterion B2. The EOO cannot be calculated due to this species being known from only one occurrence. Due to the significant threat from logging, as well as the highly restricted AOO, this species is provisionally assessed as Critically Endangered B2ab(i,ii,iii,iv,v).

We advise that targeted searches are made to refind this species and that if it is found not to be extinct, a species Conservation Action Plan is made (e.g. Couch *et al*. 2022) and that consideration is given to including this species in a protected area or demarcated Important Plant Area (as for Cameroon, e.g. Darbyshire *et al*. 2017; Murphy *et al*. 2023)

##### ETYMOLOGY

Named as a noun in apposition for Mèla (known also as Mala), 10 km north of whence the only known specimen was collected.

## Discussion

Gabon is unquestionably the centre of diversity for *Ardisia* subg. *Kamardisia* since of the ten species now known, six are endemic to the country, while only three are endemic to Cameroon, and only one is common to both countries. It is very likely that species will be found in Congo-Brazzaville, and additional species in South Region Cameroon, but it is even more likely that further species new to science will be found in Gabon itself. All the species appear to be rare and range-restricted (Peng & Cheek 2025).

The decision by Taton (1979) not to formally name his *Ardisia* sp 2, that he knew to be a distinct species, was the standard approach at that time, when it was normal to await additional material to be collected to allow a full description to be made of a species. However, today this is a luxury that we cannot afford given the march of habitat destruction. Even in countries such as Gabon where biodiversity conservation has been a priority, several conspicuous species endemic to the country appear to have become extinct, not having been refound despite a resurgence in botanical survey and collection effort (Moxon-Holt & Cheek 2021; Cheek *et al*. 2021).

## Conclusion

The majority of plant species being published as new today, three out of four, are already threatened at point of publication (Brown *et al*. 2023). This is because they often have very small ranges and so are at great risk of habitat clearance by humans. Moreover, until species have a formal scientific name, they are essentially invisible to science and a formal Red List assessment is not facilitated, making the possibility of allocation of resources to their protection remote (Cheek *et al*. 2020). This makes it urgent to characterise and formally name species before they become extinct, even if they are incompletely known, such as *Ardisia mela*.

## Acknowledgements

We especially thank Corinne Sarthou-Gasc of the curator team at Herbier National, Muséum national d’histoire Naturelle (National Museum of Natural History), Paris for superb assistance accessing specimens of the subject of this paper at P in July 2025 and also Germinal Rouhan. We also thank Mark Carine and the staff of BM, and the Africa curator-botanist team at K, led by Renata Borosova. HM thanks fellow student Caitlyn McDaniel for discussions and collaboration on *Ardisia* in 2025, during which this paper was completed as part of an MSc Project on the RBG, Kew-Queen Mary University of London Plant and Fungal Taxonomy, Diversity and Conservation.

## Data availability statement

This article has no additional data.

## References

Brown, M., Bachman, S., Nic Lughadha, E. (2023). Three in four undescribed plant species are threatened with extinction. The New Phytologist 240(4): 1340–1344. 10.1111/nph.19214

Cheek, M. & Xanthos, M. (2012). Ardisia ebo sp. nov. (Myrsinaceae), a Creeping Forest Subshrub of Cameroon and Gabon. Kew Bull. 67(2): 281–84. 10.1007/s12225-012-9362-8

Cheek, M., McDaniel, C. & Murdoch, H. (2025). Classification of Life Forms of African Ardisia (Myrsinaceae or Primulacae) (Abstract Only). – AETFAT Bulletin 50: 23.

Cheek, M., Nic Lughadha, E., Kirk, P., Lindon, H., Carretero, J., Looney, B., Douglas, B., Haelewaters, D., Gaya, E., Llewellyn, T., Ainsworth, M., Gafforov, Y., Hyde, K., Crous, P., Hughes, M., Walker, B.E., Forzza, R.C., Wong, K.M., Niskanen, T. (2020). New scientific discoveries: plants and fungi. Plants, People Planet 2: 371–388. 10.1002/ppp3.10148

Cheek, M., Tchiengué, B., van der Burgt, X. (2021). Taxonomic revision of the threatened African genus Pseudohydrosme Engl. (Araceae), with P. ebo, a new, critically endangered species from Ebo, Cameroon. PeerJ 9 :e10689 10.7717/peerj.10689

Couch, C., Molmou, D., Magassouba, S., Doumbouya, S., Diawara, M., Diallo, M. Y., Keita S.M., Koné, F., Diallo, M.C., Kourouma, S., Diallo, M.B., Keita, M.S., Oularé, A., Darbyshire, I., Gosline, G., Nic Lughadha, E., van der Burgt, X, Larridon, I. & Cheek, M. (2022). Piloting development of species conservation action plans in Guinea. Oryx, 1–10. 10.1017/s0030605322000138

Darbyshire, I., Anderson, S., Asatryan, A., Byfield, A., Cheek, M., Clubbe, C., Ghrabi, Z., Harris, T., Heatubun, C. D., Kalema, J., Magassouba, S., McCarthy, B., Milliken, W., Montmollin, B. de, Nic Lughadha, E., Onana, J. M., Saıdou, D., Sarbu, A., Shrestha, K. & Radford, E. A. (2017). Important Plant Areas: revised selection criteria for a global approach to plant conservation. Biodivers. Conserv. 26: 1767–1800. 10.1007/s10531-017-1336-6.

IUCN (2012). IUCN Red List Categories and Criteria: Version 3.1. Second Edition. – International Union for Conservation of Nature, Gland and Cambridge. http://www.iucnredlist.org/.

IUCN Standards and Petitions Committee. (2024). Guidelines for Using the IUCN Red List Categories and Criteria. Version 16. – Prepared by the Standards and Petitions Committee. https://www.iucnredlist.org/documents/RedListGuidelines.pdf

Moxon-Holt, L. and Cheek, M. (2021). Pseudohydrosme bogneri sp. nov. (Araceae), a spectacular Critically Endangered (Possibly Extinct) species from Gabon, long confused with Anchomanes nigritianus. Aroideana 44(1): 110–131. 10.1101/2021.03.25.437040

Murdoch, H. & Cheek, M. (2026). A Taxonomic Revision of the Monocaul Phanerophyte Ardisia (Primulaceae) of Gabon bioRxiv 10.64898/2026.06.26.734757

Murphy, B., Onana, J.M. van der Burgt X. M., Tchatchouang Ngansop, E., Williams, J., Tchiengué, B., Cheek, M. (2023). Important Plant Areas of Cameroon. – Royal Botanic Gardens, Kew. https://kew.iro.bl.uk/concern/books/c056b5cb-b146-4509-b98b-a7f5dd49517e

Peng, P. and Cheek M. (2025). A synoptic revision of the creeping herbaceous African species of Ardisia (Primulaceae or Myrsinaceae) with six new species from Cameroon and Gabon. Webbia 80(2) Suppl.: 139–168. 10.36253/jopt-19150

Taton, A. (1979). Contribution a l’etude du genre Ardisia Sw. (Myrsinaceae) en Afrique tropicale. Bulletin du Jardin botanique national de Belgique / Bulletin van de National Plantentuin van België 49(1/2): 81. 10.2307/3667819

Texier, N., Ngama, S., Essomba, G., Bikoukou, L., Lee, M., De Bruyne, G., Orbell, C., Ndokoua, M., Christy, P., Mipounga, H.K., Pauwels, O., Nkollo-Kema Kema, C. A.,Yombiyeni, P., Blanchet, G., Massart, A., Ikabanga, D.Paradis, A.-H. & Stévart, T. (2024). Les Zones Clés pour la Biodiversité du Gabon. Missouri Botanical Garden, USA.

